# Mosquito Mapper: a phone application to map urban mosquitoes

**DOI:** 10.1101/167486

**Authors:** Camille S.E. Guilbaud, Théophile G.D.P.V. Guilbaud

## Abstract

This paper presents mosquito mapper: an android phone application created with the goal of giving science-driven citizens the means to monitor mosquito populations in an urban environment. Mosquito mapper allows the recording of mosquito encounters as well as conditions surrounding the encounter. It also features a rudimentary identification tool. The goal of the application is to create a database and construct a map of the encounters free to consult for citizens and scientists. Such database constitutes a necessary first step for the development of useful management strategies addressing potential human health threats induced by mosquitoes. The citizen scientist may voluntarily provide other additional information on the circumstances of the encounter that may contain scientifically useful information. We describe the current features of the application, discuss their strength, limits, potential scientific value and suggest possible future extensions. The original city for which the application was developed is Berlin, Germany, but the application is coded in such a way that it is easily applicable to any urban environment.

## Introduction

Every day, more than 2.5 exabytes of digital data are generated from internet use (Gantz & Reinsel 2013). Most of these data are generated by the general public and are the primary target of companies dealing with big data. User-generated data can also be of great use for scientific purposes (Newman *et al.* 2010). A field of technologically-aided citizen science is now emerging (Silvertown 2009) which can have large scale implications for scientists, policy makers, and the citizens themselves (Dickinson, Zuckerberg & Bonter 2010; Reichman, Matthew & Schildhauer 2011). Together, we can use the large amount of data created daily to explore questions that were previously either impractical or impossible to tackle (Van Strien, Van Swaay & Termaat 2013; Cosentino *et al.* 2014). Citizen science is not a new field, but technological progress especially the development of the internet and the rapid spread of smartphones multiplies the possibilities offered by this science (Silvertown 2009; Bonney *et al.* 2014). This endeavour, however, comes with built-in limitations regarding the accuracy of the acquired data (Dickinson *et al.* 2010; Conrad & Hilchey 2011). These inaccuracies can originate from human use but also directly from the devices used for monitoring. But using the large amount of data that smartphone applications may potentially gather constitutes a promising way to temper the inherent inaccuracy that comes from dealing with untrained people using different instruments (Cohn 2008). Furthermore, citizen scientists may provide information at a scale virtually impossible to obtain from regular collection and therefore help the scientific community tackle usually difficult questions (Conrad & Hilchey 2011).

Here, we suggest using the power of citizen science to tackle the growing threats posed by mosquitoes. Mosquitoes typically constitute a nuisance in urban environments, a nuisance that may become a threat due to globalisation and climate change as diseases carried by mosquito may spread significantly faster if they enter a big centre of urbanisation (Gubler & Clark 1995; Brown *et al.* 2008). Monitoring the extent to which mosquitoes are encountered in urban environments constitutes a necessary first step for the creation of successful management (Hemme *et al.* 2010). The use of citizen science can provide a quick and cheap way of gathering such information (Cohn 2008; Bonn *et al.* 2016). Combining citizen science with the virtual ubiquity of smartphones in western urban environments may strike a winning strategy as a means of gaining information regarding the extent to which the mosquitoes cause nuisance. The trend of increased urbanisation in developed country (United Nations 2014), closely matched with market penetration of smartphone (GSM Association report 2016) suggests that such application would prove useful worldwide.

Thus, we present Mosquito Mapper, an application developed with the goal of monitoring encounters with mosquitoes in the city of Berlin. Mosquito encounters happen constantly and carry little information themselves. However, the aggregation of encounters opens the possibility to tackle questions regarding metapopulations which in turn may provide links to the spread of diseases carried by mosquitoes, or link mosquito activity within an environmental context (Cosentino *et al.* 2014). The application uses a simple touch interface to guide a user towards its different objectives. From each use of the application, there are several variables that we can access. Ultimately, such user-created database may be useful for drafting urban management policies regarding mosquitoes or be used to cast a “mosquito forecast” in the like of current weather forecast.

## 1. Alpha release

Mosquito Mapper can be downloaded on the android store (https://play.google.com/store/apps/details?id=com.sciencetogether.perecastor.mosquitomapper&hl=en). The project is currently in an alpha version. The store page contains short explanations on the goal of the application and lists which user data the application has access to. The source code for the application is accessible on GitHub (https://github.com/Layninou/MosquitoMapper).

Mosquito Mapper opens with a page displaying three buttons. A “locate” button, an “identify” button, and an information button (Figure 1). The information button simply lists the contributors to the project, acknowledges helpers and supplies contact information. The “identify” button brings a user through a short identification key. Finally, the “locate” button brings up the current user’s location followed by a short questionnaire. All information provided by a user is sent as a JSON file to a database. A user can either locate or identify the encounter, but is encouraged to do both.

**Figure.1:**
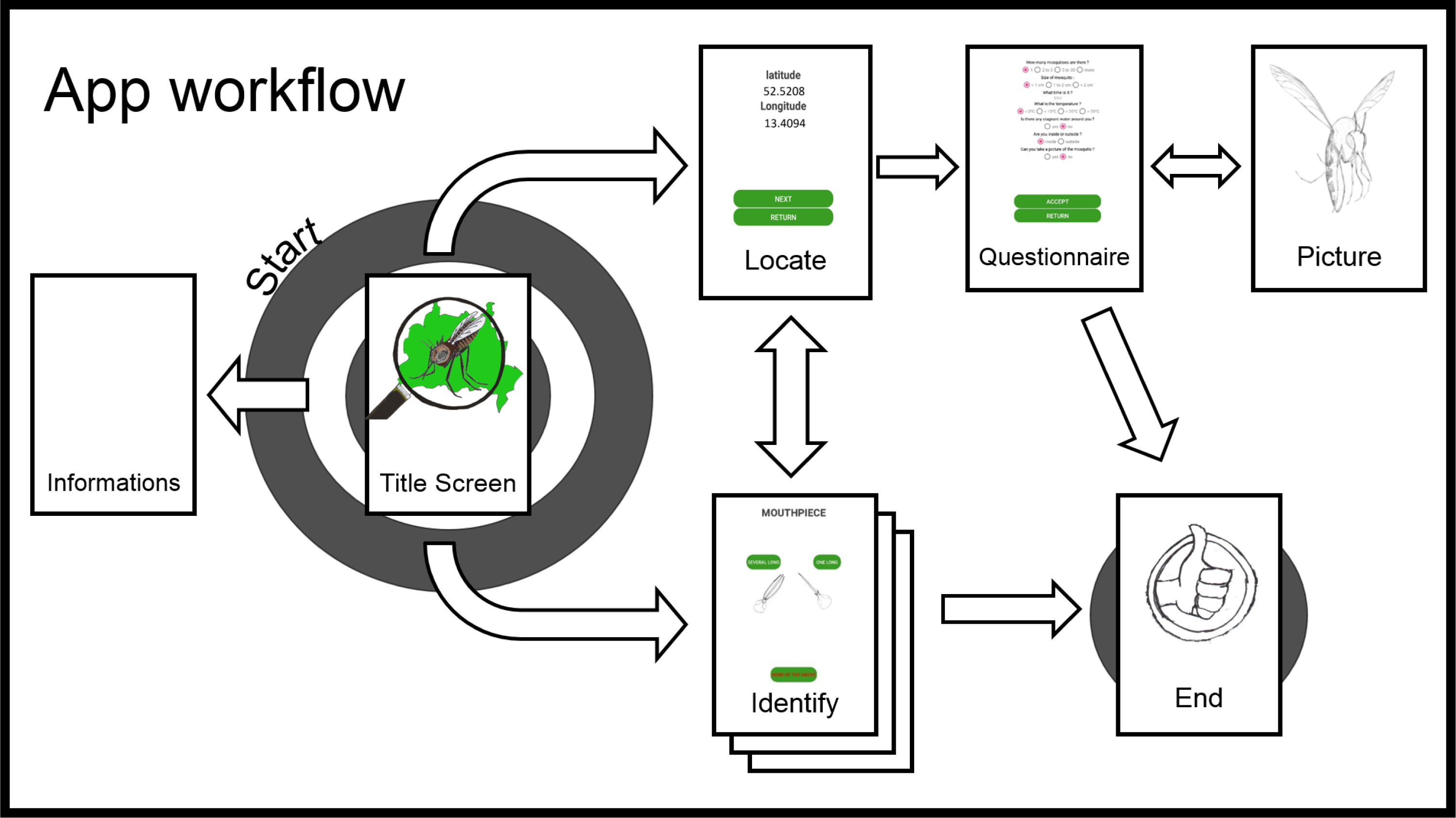
Application workflow. From the title screen, a user has three choices: starting the “locate” activity, the “identify” activity or explore the information. Information contains the name of contributors and contact information. The “locate” activity returns the user’s position followed by a short questionnaire. From there, the user may take a picture of the encountered mosquito, go to the identify activity or end the application. The “identify” activity consists of three pages with simple features to recognise on the mosquito.

The intended use of the application, that we call an “experiment” for simplicity, follows this scheme. The user opens the application, chooses whether they want to locate or identify an encounter, then follows through the chosen activity. At the end of one activity, the user is asked whether they want to follow with the other option (locate a mosquito if the user just identified one and *vice-versa*). A guide presenting the application workflow is found in Figure 1.

### Locate activity

By tapping the locate button, the users are returned their current position, then asked to proceed. The “locate” activity uses the system location service of Android. This service is standard across Android distributions and is accessed by the application upon authorization of the user. The application is therefore given access to the location device of the mobile phone (generally not the device’s GPS, the application uses various location sources) to provide the user’s coordinates. Upon starting the “locate” activity, the application creates an instance of “locate manager” and an instance of “locate listener”, the application then requests an update of the location. The “locate manager” will access the device’s coordinates (or the location of the closest Wi-Fi access or cell tower as a way to reduce the phone’s battery consumption) which is displayed to the user. The “locate listener” will track any change of position of the phone and change the displayed location accordingly. The accuracy of location provided by the application is equal to the accuracy of the mobile device and is usually precise within a range of five meters (Zandbergen & Barbeau 2011). Upon completion of this activity, the last latitude and longitude recorded is given a unique ID and sent to our server.

We decided to make this activity quick to access and finish because location is the feature we deem the most crucial. After recording the encounter, the user is asked a few questions concerning their surrounding environment and to take a picture of the mosquito. By the end of the “locate” activity, the user is asked whether they want to proceed with an identification or end the experiment.

### Identify activity

By tapping the identify button, the user is sent to a simplified identification key (Figure 2). The goal of this key is to 1) ensure that the encountered organism is a mosquito, 2) estimate which subfamily the mosquito belongs to and 3) determine whether the encountered mosquito is male or female. Upon completion of this activity, the answers given are tagged with a unique ID and sent to our server. If the locate activity has been performed before, the ID for identification is linked with the one for the location.

**Figure.2:**
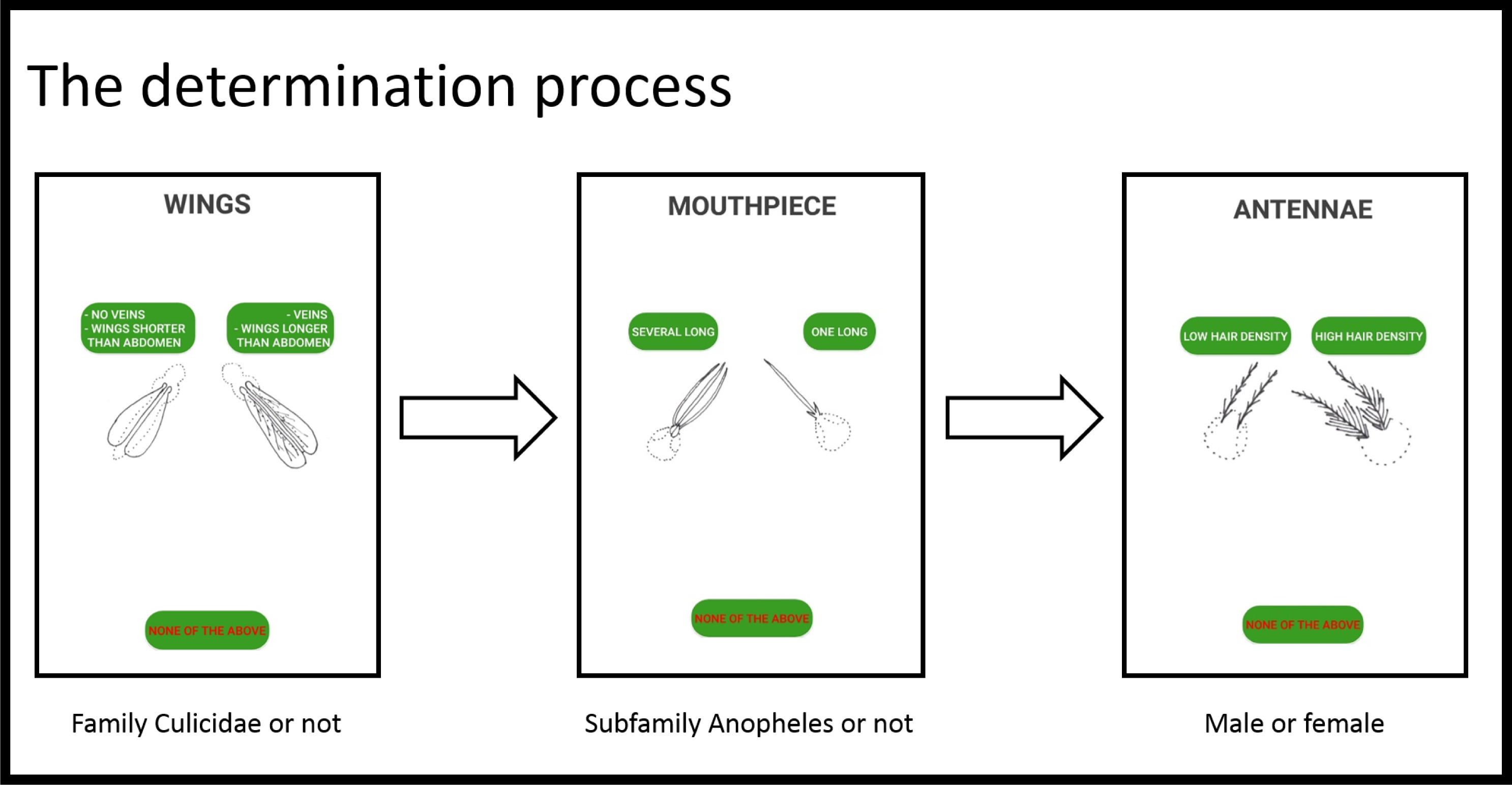
The determination process consist of three consecutive questions. The first one ensures the encountered animal is indeed a mosquito. The second one helps determining the subfamily of the encounter while the third question aims at determining the sex of the encountered mosquito.

We decided to restrict ourselves to such a low level of identification because inner city Berlin is mostly populated by mosquitoes of the genus Culex (Ann-Christin Honnen, *pers.comm*), to prevent high level of inaccuracy from the user while remaining entertaining, and to avoid asking overly complex morphological questions to untrained users (but see planned features). If the user has not completed the “locate” activity, they are asked if they want to.

### Questionnaire

After the “locate” activity, the user is asked to provide extra information regarding the context within which the encounter took place. The set of answers given is sent together with the location data (Figure 3). The questions were designed with the help of Ann-Christin Honnen from Swiss Tropical and Public Health Institute, Basel, Switzerland. The set of answers from the questionnaire will help determining the behaviour of the mosquito (e.g.: diurnal or nocturnal), their amount and the likelihood of the encounter. They are complementary to the identification activity to determine whether the encounter recorded took place with a mosquito.

**Figure.3:**
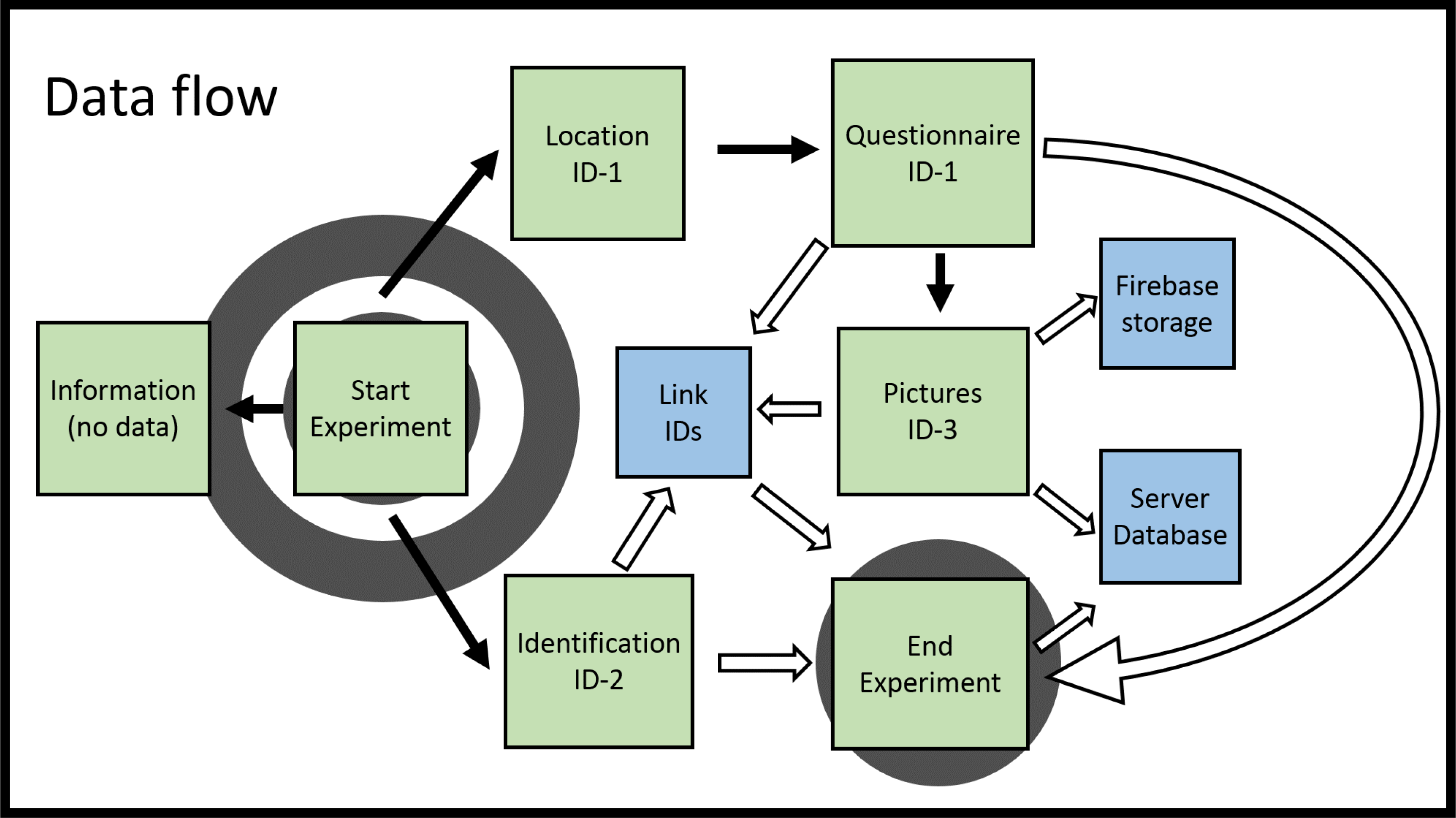
Data flow of the application. Application activities are represented in green boxes, data holders are in blue boxes. Solid block arrows represents application’s workflow while white arrows carry data. Data generated by users carry unique IDs and are linked before being sent to the server. Activities performed alone are not linked. Picture data are sent in two places: a Firebase storage where they can be viewed directly and the server where they are stored as a byte array.

### Pictures

At the end of the questionnaire, the user is asked to take a picture of the mosquito encountered. By choosing yes, the phone camera opens. Ideally, the pictures taken by citizens could be used later by zoologists as a way to improve species distribution maps although preliminary tests were inconclusive. Pictures are usually large files, which could clutter our database. To reduce the size of images, these are modified locally using a BLOB (Binary Large Object) procedure. This procedure transforms the bitmap image taken with the camera into a bytes array. The bytes array is then sent directly to the server. Image quality is decreased to a quarter of the original picture.

### End of an Experiment

Upon completion of any of the two activities, the user is directed to a last page where they are asked to send all information. When the user taps the “send experiment” button, all data associated with the unique experiment ID is confirmed in our database. Data are stored as JSON objects (JavaScript Object Notation) divided in three branches. One for the questionnaire and location data, one for the identification data and one for the pictures.

### Miscellaneous

With this alpha release, all information and pictures are sent to a private server, by downloading the application, users agree to provide us with the data and give us the right to store it for scientific purposes. The type of data collected and the moment they are collected is detailed in Figure 3. They do not contain any personal information. Ultimately, our goal is to make all the data collected freely accessible on a website currently under construction.

## 2. Strengths and limits

Developing a smartphone application for citizens and scientists to use comes with a suite of strengths and limits that depends to a large degree on variables that are unfortunately not in our control. First and foremost, the amount of data generated by the application will depend on the number of users. The usefulness of data will also depend on the users. A critical goal of our application is the ability to grossly assess the population dynamic of mosquitoes through time. This requires a regular feed of data. One downside of our application then becomes the possibility of having large amounts of data but distributed in a way that prevents their use for certain types of investigation. For example, it is possible that many data are collected at the end of spring when mosquitoes emerge and very little data afterwards. We will include regular notifications in the final release (see planned features) as a way to mitigate this issue. In its present state, the application relies on the willingness of participants to provide data as well as the introduction of new users.

There is an inherent difference between the existing data accounted for by the user and the true temporal and spatial distribution of mosquitoes. Such inaccuracies in typical experiments are overcome through the use of dedicated statistical methods (Fithian *et al.* 2015). The accuracy of our data will strongly depend on the frequency and the amount of recorded encounters.

The main use of our application will probably remain limited to the city of Berlin were we have the capacity to advertise our work on a regular basis to an interested audience. The use of Mosquito Mapper on larger scale, however useful it may be, would rely on time investment from other people.

Developing a smartphone application for big data seems to come with a lot of uncertainties. Before such endeavour is widely adopted, there is a great possibility that our work produces little useful studies. However, because such application can remain dormant at little to no cost, we believe it offers scientists a great opportunity to study mosquitoes in a way that can be both cheap and efficient. In fact, the strengths and limits described above all rely on the amount of users and their willingness to provide accurate and regular data. All limits eventually turn into strength once a critical amount of users is reached. As such, the value of an application such as Mosquito Mapper can only increase with time, as even low amount of data generated per unit of time will compound into appreciable quantities.

## 3. Value of the data

For the scientist, the data collected through the application could be used for several purposes. On the one hand, GPS data together with the number of recorded encounters plus their time frame can be used to generate dynamic maps of mosquito populations. On the other hand, these data can be used to predict changes of mosquito populations’ sizes for modelling purposes. The pictures received could serve as a repository that anyone could use to create species distribution maps or compare phenotypes within the same species. As time passes, the potential from the data obtained increases. And each of the scientific experiments described above can be put into a temporal framework. Finally, as technology improves, there might be new ways of studying the data we aim at collecting or ways that already exist but that we have not thought of. The possibility of serendipitous discovery is the main reason we plan to offer free access to all the data collected for anyone.

For the citizen, there are two main positive outcomes of the existence of the application. There is first the active part where the citizens are engaged in an entertaining activity. Hopefully, the data collectors will have fun using the application. The citizen scientist, actively participating in the construction of a database they can profit from, should feel more closely connected to the scientific community. On the passive side, the predictive maps of mosquito presence could serve as an information tool at the same level as the weather forecast, so that people are informed where not to go if they want to minimise the probability of a mosquito encounter. Finally, the data could be used by municipalities to initiate public health actions in order to reduce mosquito presence.

## 4. Planned features

The alpha release of the application is meant to showcase the possibilities of citizen-driven data collection with the help of smartphones. By trying to make the data collection as easy and fast as possible, our aim is to allow children to take part. As a result, we wish to add features to make the application more “fun” to use. One way to do this is to gamify the application which means adding game-like features that would make using the application more engaging. We envision this process as follows: each use of the application would generate experience points that would lead the user to change levels. We will then add a leader board where users could compare their performance to others. Another direction for gamification will be the addition of badges for certain behaviour such as identifying a certain amount of mosquitoes or with a certain regularity. Developing the application to be more game-like implies various changes to the way the database is currently structured that would make the application more lightweight but would also consume more mobile data.

A big issue of the program is that it consists of a presence-only dataset that is known to suffer from bias (Blossey & Hunt-Joshi 2003; Fithian *et al.* 2015). Presence-absence datasets are more reliable. Therefore, we will incorporate notifications in the application to encourage users to report places and times when no encounter took place. The downside of such feature is that it may become annoying. The notification shall therefore only trigger rarely (once a week), with the possibility to turn the feature off.

Ultimately, we wish to include more precise determination keys. The ability to use more complex keys may be limited to users reaching a certain “level”, thus ensuring their willingness to participate fairly. The determination of mosquito species may be difficult and may vary wildly depending on geographical location. As a result, such feature may be limited to a handful of countries.

## Acknowledgments

C.S.E Guilbaud is funded by the Swiss National Funds (grant number P2ZHP3_158949). C.S.E Guilbaud is thankful to B.Tietjen, J.Petermann and AC. Honnen for their help and support during the writing of the manuscript. The authors declare no competing interests.

## Data availability statement

All non-social data (i.e. location and pictures) will be accessible on a website currently under construction. In the meanwhile, data is accessible through Github (https://github.com/Layninou/ScienceTogether).

